# Modelling selection response in plant breeding programs using crop models as mechanistic gene-to-phenotype (CGM-G2P) multi-trait link functions

**DOI:** 10.1101/2020.10.13.338301

**Authors:** M Cooper, O Powell, KP Voss-Fels, CD Messina, C Gho, DW Podlich, F Technow, SC Chapman, CA Beveridge, D Ortiz-Barientos, GL Hammer

**Author notes:** Corresponding Author: Mark Cooper.

## Abstract

Plant breeding programs are designed and operated over multiple cycles to systematically change the genetic makeup of plants to achieve improved trait performance for a Target Population of Environments (TPE). Within each cycle, selection applied to the standing genetic variation within a structured reference population of genotypes (RPG) is the primary mechanism by which breeding programs make the desired genetic changes. Selection operates to change the frequencies of the alleles of the genes controlling trait variation within the RPG. The structure of the RPG and the TPE has important implications for the design of optimal breeding strategies. The breeder’s equation, together with the quantitative genetic theory behind the equation, informs many of the principles for design of breeding programs. The breeder’s equation can take many forms depending on the details of the breeding strategy. Through the genetic changes achieved by selection, the cultivated varieties of crops (cultivars) are improved for use in agriculture. From a breeding perspective, selection for specific trait combinations requires a quantitative link between the effects of the alleles of the genes impacted by selection and the trait phenotypes of plants and their breeding value. This gene-to-phenotype link function provides the G2P map for one to many traits. For complex traits controlled by many genes, the infinitesimal model for trait genetic variation is the dominant G2P model of quantitative genetics. Here we consider motivations and potential benefits of using the hierarchical structure of crop models as CGM-G2P trait link functions in combination with the infinitesimal model for the design and optimisation of selection in breeding programs.

## Introduction

Plant breeding programs are designed to develop improved cultivated varieties (cultivars) of crops for use in agriculture. They have a long history and have served an important role in improving crop productivity (Allard 1960, 1999, Wricke and Weber 1986, Fehr 1987a,b, Hallauer and Miranda 1988, Blum 1988, Cooper and Hammer 1996, Bernardo 2002, Duvick et al. 2004, Fischer et al. 2014, Smith et al. 2014, Hammer et al. 2019, Voss-Fels et al. 2019a). Through the iterative cycles of breeding programs, plant breeders utilise the accessible genetic variation for traits, available through elite germplasm and other genetic resources, to improve the genetics of multiple traits of crop cultivars (Figure 1, Table 1). Today many technologies can be applied to change the genetic content of plants and discover new pathways for trait improvements (Tester and Langridge 2010, Morrell et al. 2012, Wallace et al. 2018, Bailey-Serres et al. 2019). Here we will focus our considerations on the genetic improvement of traits by directional selection within the context of plant breeding programs (Figure 1; Cooper et al. 2014, Walsh and Lynch 2018). Selection for cultivar performance and breeding value has been the foundation for long-term genetic improvement of crops (Duvick et al. 2004, Smith et al. 2014, Walsh and Lynch 2018, Voss-Fels et al. 2019a). For purposes of discussion, crop grain yield will be considered as the ultimate trait of interest (Evans 1993, Fischer et al. 2014). Importantly, grain yield is a multi-trait outcome of crop growth and development processes and responses to diverse environmental conditions (Evans 1993, Cooper and Hammer 1996, Messina et al. 2009, Connor et al. 2011, Fischer et al. 2014). Within this context we argue that the crop sciences (Messina et al. 2020) together with advances in plant and crop models have the potential for important new roles in improving the design and effective operation of breeding programs within the context of the future needs for crop improvement (Hammer et al. 2019, 2020). To realise these opportunities plant and crop models will have to be designed to take advantage of advances in understanding of trait genetic architecture and the principles of quantitative genetics (Cooper et al. 2002a,b, 2005, 2009, Hammer et al. 2006, 2019). Here we introduce a quantitative genetics perspective of approaches for linking trait genetic models with mechanistic crop models to enhance our understanding of the genetic architecture of complex traits, such as grain yield of crops, and to explore new prediction applications for breeding.

**Table 1.** Definitions of common terms represented in Figure 1 and their uses for plant breeding and quantitative genetics.

| Term | Definition and use |
| --- | --- |
| Germplasm | A collective term used in plant breeding to refer to the complete set of genetic resources that can be accessed and used within a plant breeding program. Elite germplasm is often used to identify the set of germplasm that has been improved by the breeding program and represents the genetic diversity that is actively being used by the plant breeder. |
| Reference Population of Genotypes: RPG | The population of germplasm as developed by a breeding program that is in use as the reference set of genotypes and for which any estimates of genetic variation and gene effects will apply. The classical reference population of genotypes (RPG) for population and quantitative genetics is a population in Hardy-Weinberg equilibrium. Hardy-Weinberg equilibrium applies when the population is large and maintained by random mating of individuals. There are many other mating strategies applied in plant breeding, involving different combinations of cross-pollination (crossing) and self-pollination (selfing). These diverse mating structures frequently result in populations of genotypes that diverge from the expectations for the classical Hardy-Weinberg random mating population structure. Over multiple cycles of the breeding program, complex pedigree relationships are formed connecting individuals and creating the structured RPGs that are the targets for selection in breeding programs. An understanding of the structure of the RPG is required to correctly interpret and use estimates of genetic variation and genetic effects for traits within a breeding program. |
| Genotype | Theoretical reference to an individual sampled or selected from a reference population of genotypes (RPG). Genetic information for the individual can be defined for one or more genes or a whole genome. For a single gene, the genotype refers to the allele combinations for a particular gene. For a gene with two alleles, **A** and **a** there are three possible genotypes, i.e. the two homozygotes **AA** and **aa** and one heterozygote **Aa**. Genetic variation can be quantified at the allele and genotype levels for samples of individuals from the RPG. In experimental studies, entries are often referred to as “genotypes”. We recommend avoiding this use of the term genotype whenever there is no characterisation of the experimental entries based on genetic information or connection of the entries to a defined RPG. For such cases, we recommend alternative references to the individuals as entries in experiments as defined below. |
| Phenotype | The observable measurement of a trait value for an individual or collection of individuals. The phenotypic value for a trait observed in an experimental situation is considered to be an outcome from the combination of genotype and environmental effects and their interaction. |
| Individual | Experimental complement to the theoretical reference to a genotype. Individual is used to refer to an identifiable entry that can be studied experimentally: e.g., experimental line, cultivar. This is recommended to avoid any incorrect referencing of an entry in an experiment as a genotype for situations where genetic information is not directly investigated and where there is no clear connection of the sample of individuals to a reference population of genotypes. |
| Trait Genetic Architecture | Trait genetic architecture is a quantitative genetics construct/term/framework/expression. For a defined trait, genetic architecture is used to refer to the set of genes and their alleles that determines genetic variation for the trait within the context of a reference population of genotypes (RPG). Commonly considered components include; the number of genes and their positions within the genome, the number of alleles for a gene and their frequencies within the RPG, the effects of the genes and alleles on the genotypic values of individuals and genetic variation among the individuals within the RPG. The total genotypic value can be partitioned into within and among gene effects. The within gene partition is among the alleles of a gene, interpreted in terms of additive and dominance components. The among gene effects is among allele combinations for different genes, interpreted in terms of components of epistasis. The framework for the partitioning of genotypic value can also be used to investigate interactions with different environments. |
| Average Effect of Allele Substitution | The average effect of an allele and an allele substitution are quantitative genetics concepts. The average effect of an allele is the mean deviation from the population mean of individuals that received the allele of interest from one parent, with the allele received from the other parent having come at random from the RPG (Falconer and Mackay, 1996). For a graphical example see Figure 3b. Note that the definition of the average effect is dependent on mating structure of the population, i.e. it strictly applies to the RPG (Falconer, 1985). Thus, the “average effect” of all the possible “allele substitutions” within the RPG is dependent on the allele and genotype frequencies within the RPG in combination with the differences in trait values between the genotypes (Figure 3b). In the context of a breeding program, the average effect gives an estimate of how much the average trait value of the RPG could be improved when a favourable allele is increased in its’ frequency, e.g. through targeted crossing. With increasing frequency of the favourable allele in the RPG, the size of its’ average effect diminishes and ultimately becomes zero once it gets fixed, i.e. once all individuals in the RPG carry the favourable allele. The average effect of allele substitution is an important determinant of the additive genetic variation and variation for breeding values within the RPG (Falconer and Mackay, 1996). |
| Additive Genetic Variance | The additive genetic variance is a measure of the component of the total genetic variance that is associated with the average effects of the alleles of genes within the reference population of genotypes. This additive component of variance measures the component of the total genetic variance directly transferred from parents to progeny through sexual reproduction, through transmission of alleles from parents to progeny. The additive genetic variation is the target of the selection predictions made using different forms of the breeder’s equation. |
| Breeding Value | A quantitative genetics statistic that provides a measure of the consistency with which the trait values of an individual can be transferred through sexual reproduction to their progeny. The breeding value of an individual is determined by the average effects of the alleles that are possessed by the individual within the context of the reference population of genotypes. This should be considered in terms of the summation of the effects across all genes involved in the trait genetic architecture. The variance among individuals for their breeding values determines the additive genetic variance and narrow sense heritability for the trait within the reference population of genotypes. |
| Heritability | A quantitative genetics statistic that measures the ratio of genetic variance to phenotypic variance for a reference population of genotypes. The ratio can be defined in many different ways. A common distinction is between broad sense heritability and narrow sense heritability. Broad sense heritability measures the ratio of total genetic variance to phenotypic variance. Narrow sense heritability measures the ratio of additive genetic variance to phenotypic variance. As such narrow sense heritability is the appropriate form of heritability for use in the breeder's equation for prediction of response to selection when sexual reproduction is involved. The details of the breeding strategy require attention to determine the correct form of the narrow sense of heritability for prediction. Another important consideration for definition of heritability is the unit of observation for trait measurement and unit of selection. |
| Target Population of Environments: TPE | The Target Population of Environments for a breeding program refers to the complete set of environmental conditions that can occur within the target on-farm conditions of the breeding program. This could extend from the environmental conditions encountered within a small regional scale to a multi-continental scale depending on the breeding program. The level of resolution of the different environmental-types defined by environmental conditions depends on the level of resolution that is possible when measuring important biophysical environmental variables. Many levels of resolution are possible providing a range of coarse-grained to fine-grained definitions of the environment-type composition of the target population of environments. Clear definition of target environment-types is required for any investigations and breeding programs that encounter strong influences of Genotype-by-Environment (GxE) interactions. Measurement of environmental variables that can be used to distinguish environment-types provides a basis for construction of predictions that can take into account GxE interactions. Agronomic management is considered as a special category of the environmental component of the target population of environments. Agronomic management provides a control variable through which farmers can account for environmental variation that impacts on-farm productivity. Plant breeders and crop agronomists consider the importance of Genotype-by-Environment-by-Management (GxExM) interactions. |
| Multi-Environment Trials: METs | Multi-Environment Trials are the experimental component of breeding programs. Samples of genotypes from the RPG are evaluated in samples of environments from the TPE. The samples of environments can include a diverse range of situations from specially designed controlled environment facilities to farmer's fields using the management practices of the farmers. The correct design and analysis of METs to reliably represent the RPG and TPE is foundational to design of model-based prediction methods for plant breeding. |
| Envirotyping | A collection of proximal and remote sensor-based technologies, methodologies and analysis tools that have been developed to characterise biophysical properties of environments and enable their definition as environment-types and measure their frequency of occurrence for the TPE. The same methods can be applied to characterise the environments sampled in METs to provide measures of how well the samples of environments realised in any MET represents the TPE. The environmental characterisation measurements and the classification of environments as environment-types can be applied separately or in combination in statistical models to construct predictors for applications in plant breeding and to model plant breeding programs to account for GxE and GxExM interactions. |

**Figure 1.**
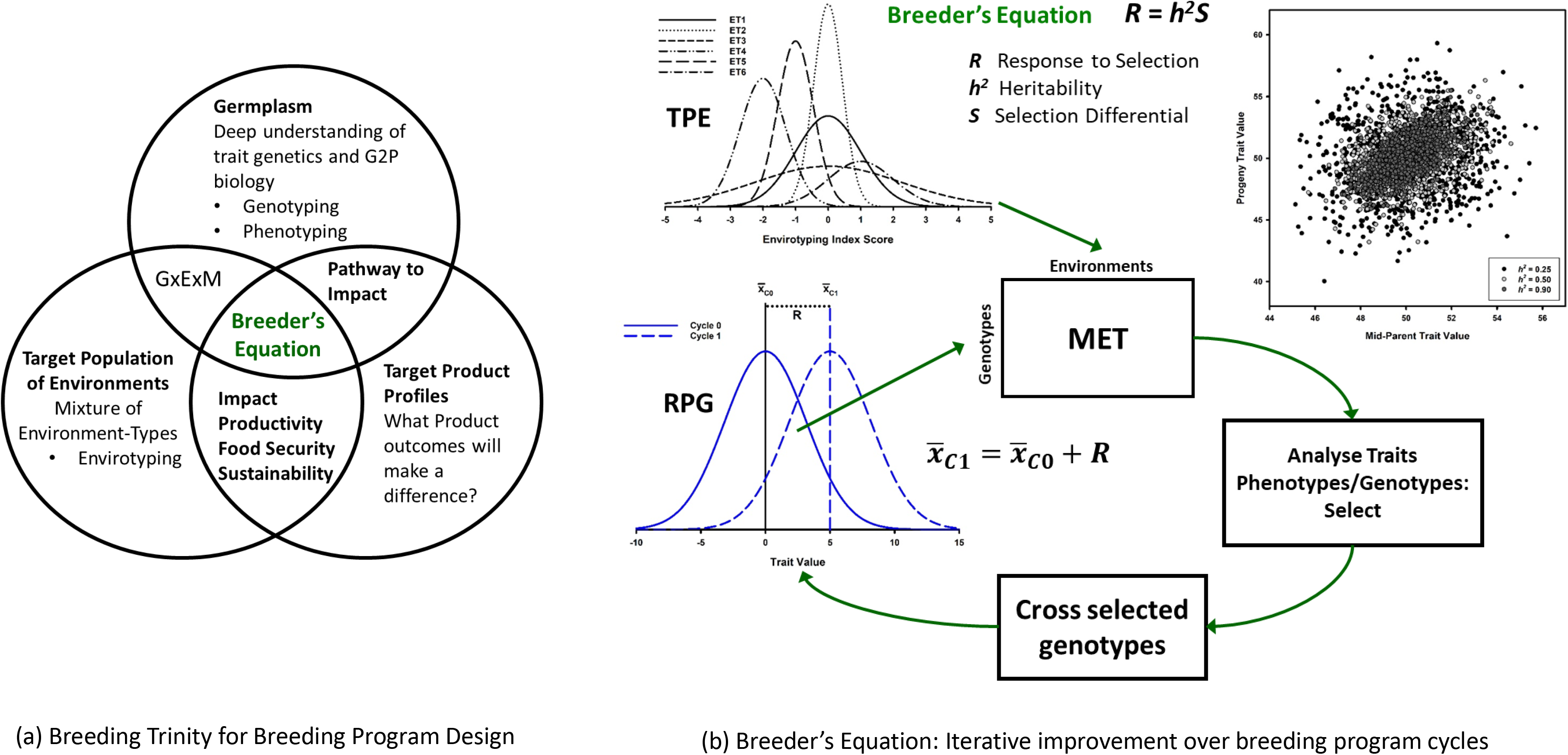
Schematic representation of key steps involved in a breeding program cycle. The example depicts one cycle of selection. The selection is applied to the Reference Population of Genotypes (RPG), beginning with Cycle 0 (C0) to create Cycle 1 (C1). Individuals (genotypes) are sampled from C0. The C0 individuals are tested in a Multi-Environment Trial (MET), based on a sample of environments taken to represent the mixture of environment-types within the Target Population of Environments (TPE). The sample of environments obtained for a MET can be characterised by measuring environmental variables to determine the environment-types sampled. Trait phenotypes are measured on the individuals within the sample of environments obtained in the MET and analysed. The genotypes of the individuals can also be measured as molecular marker fingerprints. Marker-trait associations can be established and used as a basis for selection decisions. Based on the results of the analyses, the C0 individuals are sorted into a select or a reject group. The selected individuals are retained and used as parents in a planned crossing scheme to create a new generation of progeny to establish C1. The heritability (*h*^*2*^) parameter of the breeder’s equation defines the expected predictive relationship between the trait values of the parental individuals of C0 and the trait values of their progeny (C1) (the insert shows examples of expected mid-parent and progeny associations for three levels of trait heritability; *h*^*2*^ = 0.25, 0.50, 0.90). Through the crossing step of the breeding program cycle the alleles of the genes that determine the trait phenotypes of the selected individuals from C0 are transmitted to the progeny of the next generation. The progeny become the C1 individuals of the next cycle of the RPG. New genotypes are created in C1. The breeder’s equation can be applied to the results of the MET to predict the expected genetic improvement (*R*) of the RPG trait mean between C0 and C1. Numerical values included in the schematic were simulated for a genetic model based on 100 genes, each with two alleles. Table 1 provides a definition and explanation of uses of common plant breeding terminology used in the schematic.

While we focus on plant breeding programs, it is understood that they do not operate in isolation to achieve sustainable improvements in on-farm crop productivity. Successful crop improvement programs combine the genetic improvement outcomes from breeding programs with the recommendations from agronomy research to deliver on-farm improvements in crop productivity (e.g., Duvick et al. 2004, Hammer et al. 2014, Fischer et al. 2014, Messina et al. 2020, Cooper et al. 2020). Thus, breeding programs and agronomy research programs are strongly interdependent. However, within the dominant crop improvement paradigm of today they operate sequentially, typically with limited or loosely coupled levels of integration. Within this sequential approach, breeding programs first develop new crop cultivars, arguably with limited attention to sampling the full range of agronomic management possibilities used by farmers. Then agronomic research programs follow, focussing on the development and optimisation of crop management strategies for the new cultivars. Both plant breeders and agronomists are interested in traits and how they contribute to on-farm crop productivity through improved yield, yield stability, grain nutrition for human and animal consumption and grain quality for many commercial uses. However, they differ in the types of models they use to study traits for such common objectives. We consider how mechanistic crop models can be developed to connect the modelling objectives of plant breeding and quantitative genetics with those of crop agronomy for important traits contributing to grain yield potential and yield stability (Cooper et al. 2009, Hammer et al. 2019, Messina et al. 2020).

Mechanistic crop models have a long history of development (Holzworth et al. 2014, Jones et al. 2016) and have been used for many applications to support agronomy research. However, in contrast, for plant breeding there has been much less consideration of the potential applications of mechanistic crop models to study traits. Nevertheless, given the long-term nature of breeding programs, there has been parallel interest in applications of simulation methods for modelling plant breeding programs (Podlich and Cooper 1998, Cooper et al. 2002a,b, Li et al. 2012, Bernardo 2020). Here, the statistical gene-to-phenotype (G2P) models of quantitative genetics are used to study and represent the genetic architecture of traits and the effects of genes on trait variation (Falconer and Mackay 1996, Lynch and Walsh 1998, Walsh and Lynch 2018). However, to date these quantitative genetic models of trait genetic architecture have received little attention in agronomy research and in the development of mechanistic crop models. These differences in trait modelling approaches can create barriers to their integration for accelerated crop improvement. However, understanding their potential connections creates new opportunities (e.g., Cooper et al. 2002a,b, 2005, 2016, Chapman et al. 2002, 2003, Hammer et al. 2006, 2019, Chenu et al. 2009, 2017, 2018, Messina et al. 2011, 2018, Technow et al. 2015, Onogi et al. 2016, Bustos-Korts et al. 2019a,b, Peng et al. 2020). Here we provide an overview of the progression from the trait G2P models of quantitative genetics to applications using mechanistic crop models (CGM-G2P). The review is orientated from a perspective of seeking opportunities to use crop models with quantitative genetics to integrate plant breeding and agronomy to enhance prediction of crop improvement outcomes.

The overarching objective of the manuscript is to demonstrate how an integrated crop improvement strategy, based on trait genetics, crop physiology, breeding and agronomy, can be enabled if we can use CGM-G2P multi-trait link functions within the framework of quantitative genetics for design and operation of plant breeding programs. The manuscript provides: (1) an introduction to the breeder’s equation and the infinitesimal G2P model of trait genetic architecture that are widely applied in quantitative genetic theory to model plant breeding programs, (2) possible extensions of the infinitesimal model of quantitative genetics using the hierarchical structure of crop growth models (CGM-G2P), (3) demonstration of applications of the hierarchical CGM-G2P models to plant breeding, and (4) discussion of the implications CGM-G2P models for plant breeding.

## Modelling Plant Breeding Programs

The objectives of breeding programs are defined in terms of trait targets for genetic improvement (Figure 1a). The targets are based on the required combinations of plant traits for superior performance of a new cultivar and also the current levels of the traits possessed by the cultivars that are to be replaced by the new products of breeding programs (e.g., Hallauer and Miranda 1988, Fehr 1987a,b, Bernardo 2002). The trait targets for a new cultivar may be defined in terms of specific levels of expression of the trait phenotype. For example, a specific threshold level of disease resistance, drought tolerance and grain quality may be required for a cultivar to be useful for the production systems of farmers. Alternatively, the targets can be defined in terms of levels of trait phenotypes that are superior to those of the dominant cultivars currently in use by farmers. Combinations of both approaches are common; e.g. create cultivars with new combinations of disease resistance genes, 5% improved grain yield under defined drought conditions and 5% reduced post-harvest storage and processing costs compared to a given cultivar. To achieve these objectives plant breeders design and operate breeding programs over multiple cycles (Figure 1b). The breeding programs use selection, in combination with segregation and recombination of the alleles of the genes controlling the traits, to create the targeted change in genetic control of traits. The genetic basis of the change enabled by selection is achieved by increasing the frequencies of favourable alleles and creating new combinations of the favourable alleles across all of the genes that contribute to the control of variation for the target traits and which are polymorphic within the reference population of genotypes (RPG). The genetic changes required to move from the current genotypes to the improved target genotypes can be modelled in terms of genetic trajectories in multi-dimensional G2P space (Podlich and Cooper 1998, Gavrilets 2004, Walsh and Lynch 2018). Based on selection theory, a genetic trajectory is achieved by changing the frequencies of the favourable alleles of the genes within the RPG, which in turn enables the creation of new genotypes through assembling new combinations of the alleles across many genes (e.g., Falconer and Mackay 1996, Podlich and Cooper 1998, Hammer et al. 2006, Messina et al. 2011, Walsh and Lynch 2018). Depending on the trait and structure of the RPG, the numbers of genes, or Quantitative Trait Loci (QTL), involved in the genetic architecture of a trait have been estimated to range from few to many hundreds (e.g., Barton and Keightley 2002, Cooper et al. 2005, Boer et al. 2007, Buckler et al. 2009, Mace et al. 2019, Wang et al. 2020). Quantitative genetics provides the theoretical framework and methods for modelling the genetic trajectories that underlie the genetic improvement enabled through a breeding program (Falconer and Mackay 1996, Lynch and Walsh 1998, Walsh and Lynch 2018, Wisser et al. 2019). Thus, to model breeding programs it is necessary to be able to model trait genetic architecture, trait G2P relationships, and how selection brings about genetic change for traits within the context of the structured RPG of the breeding program.

To assess contributions of allele effects to the selection response for a trait in breeding applications requires consideration of the breeder’s equation in combination with three breeding concepts (Figure 1, Table 1): Germplasm, the Target Population of Environments (TPE), and Trait Product Profiles. Germplasm represents the structured pools of genetic resources that are available to the breeder, and the organisation of the genetic diversity available through the germplasm (standing genetic diversity) into the RPG used in the breeding program. The TPE represents the mixture of environment-types for which cultivar performance is evaluated and targeted. Trait Product Profiles represent the important trait targets required by cultivars to achieve improved performance within the TPE. This “breeding trinity” sets the scene for the operation of the breeding program. Together the germplasm and the TPE determine the biophysical properties and genotype-by-environment-by-management (GxExM) context of the agricultural system within which the breeding program operates (Messina et al. 2009, Cooper et al. 2020). The Trait Product Profiles identify the targets for genetic improvement of crops. The breeder’s equation quantifies the speed with which the Trait Product Profiles can be achieved by the breeding program using selection applied to the standing genetic variation that is accessible to the breeder within the RPG.

New genotypes are created over breeding program cycles by manipulating, selecting and recombining trait genetic variation within the context of the genetic diversity for a RPG (Figure 1b). Trait performance for the new genotypes is measured and evaluated within the context of the range of environmental conditions expected for a TPE (Figure 1b). Breeding programs are conducted for multiple cycles (Figure 1b). Each cycle produces a new cohort of cultivars. The Target Product Profiles are rarely achieved in one cycle of a breeding program. Thus, the products of the breeding program cycles provide a continuous sequence of new cultivars with progressively improving trait performance for the TPE, moving towards the Target Product Profiles (e.g., Duvick et al. 2004, Smith et al. 2014, Atlin et al. 2017, Voss-Fels et al. 2019a). When the new cultivars are widely grown by farmers throughout the TPE they can contribute to improved crop productivity (Fischer et al. 2014, Atlin et al. 2017, McFadden et al. 2019). We also note that the Target Product Profiles of a breeding program are rarely static. They change as the conditions of agricultural systems and needs of society change.

## Plant Breeding and Quantitative Genetics

The methods and technologies for design and conduct of plant breeding programs are continually evolving (e.g., Paterson et al. 1988, Lande and Thompson 1990, Podlich and Cooper 1998, Meuwissen et al. 2001, Cooper et al. 2014, Bevan et al. 2017, Watson et al. 2018, Voss-Fels 2019b, Ramstein et al. 2019, Bernardo 2020, Reynolds et al. 2020). However, since the first half of the twentieth century, quantitative genetics has provided the dominant theoretical framework for studying trait genetic architecture and trait genetic variation within the RPG of breeding programs (e.g., Fisher 1930, Kempthorne 1957, Falconer 1960, Falconer and Mackay 1996, Lynch and Walsh 1998, Walsh and Lynch 2018). Plant breeders have developed and applied quantitative genetic theory to guide the design of plant breeding programs (e.g., Hanson and Robinson 1963, Wricke and Weber 1986, Hallauer and Miranda 1988, Nyquist and Barker 1991, Comstock 1996, Bernardo 2002, Holland et al. 2003). Within quantitative genetic theory, the breeder’s equation (Figure 1b) provides a foundation for application of quantitative genetics selection theory to plant breeding and to populations in general (Walsh and Lynch 2018). The breeder’s equation also provides a framework for considering how to extend trait G2P models to incorporate the advances in understanding of trait physiology and plant responses to environmental conditions as they are quantified within crop models. Below, we develop further this quantitative connection between trait genetics and physiology through crop models.

## The breeder’s equation as a framework for linking genetics to crop growth models

The breeder’s equation provides a modelling framework, grounded in quantitative genetics theory, for predicting response to selection. Therefore, the breeder’s equation has been used to design and optimise breeding programs for accelerated genetic improvement of traits by selection (Figure 1). As we will discuss, it also provides a framework for modelling G2P relationships for traits and as such, provides a foundation for linking genetics to crop growth models. The basic form of the breeder’s equation *R* = *h*^*2*^*S* predicts trait response (*R*) from a cycle of selection as the product of the target trait heritability (*h*^*2*^) and the selection differential (*S*) applied to the phenotypic variation for the target trait within the RPG. In this form, the breeder’s equation emphasises the importance of enhanced trait phenotyping to achieve high trait heritability relevant to the RPG as a pathway to increase response to selection (e.g., Araus and Cairns 2014, Araus et al. 2018, van Eeuwijk et al. 2019, Reynolds et al. 2020). Extensions enable further considerations of trait genetics and opportunities to increase response to selection (e.g., Nyquist and Baker 1991, Comstock 1996, Holland et al. 2003, Walsh and Lynch 2018). One useful form of the breeder’s equation expresses trait variation in terms of the trait phenotypic variation. The standardised selection differential (*i*) defines the selection differential in terms of the standard deviation of the trait phenotypic variance within the RPG, *i* = *S*/*σ*_*p*_. By substitution, the basic form of the breeder’s equation can now be written as *R* = *ih*^*2*^*σ*_*p*_. This extended form applies to many situations where selection decisions are based on the observed variation for trait phenotypes (Nyquist and Baker 1991, Holland et al. 2003). The development of genomic prediction methods for breeding applications has enabled selection based on trait genetic variation measured at the genome sequence level (Muewissen et al. 2001, Bernardo and Yu 2007). For this, we require another form of the breeder’s equation. By expressing the trait heritability as the ratio of additive genetic variance to phenotypic variance (*h*^*2*^ = *σ*^*2*^_*a*_/*σ*^*2*^_*p*_) we can substitute and simplify to obtain a form of the breeder’s equation *R* = *ir*_*a*_*σ*_*a*_. Here *r*_*a*_ is a measure of predictive accuracy, based on the correlation between trait breeding values and phenotypic values, and *σ*_*a*_ is the square root of the trait additive genetic variance for the RPG. Thus, response to selection is predicted as the product of the intensity of selection, the accuracy of prediction that can be achieved and the magnitude of variation within the RPG for trait breeding values of the selection units. This third form of the breeder’s equation emphasises two issues relevant to developing trait G2P models. Firstly, understanding the magnitude and form of trait additive genetic variation for the target traits within the RPG. Secondly, the accuracy with which genomic information can be used to predict the additive genetic contributions of the standing trait genetic variation to the breeding value of the individuals selected and used as parents to create the RPG for the next cycle of the breeding program. To understand and illustrate the opportunities for enhanced prediction and new breeding methods that are created by linking trait genetics with crop growth models we need to unpack the breeder’s equation from the perspective of trait G2P link functions to provide a foundation for developing and discussing the extended CGM-G2P link functions. The breeding program as presented in the schematic in Figure 1, together with this framework of the breeder’s equation, motivates opportunities to link trait genetics with crop models (Voss-Fels et al. 2019b).

## Motivations for linking quantitative genetics and crop modelling

Genetic improvement of complex traits, such as grain yield, is a long-term process requiring many cycles of a breeding program (e.g., Duvick et al. 2004, Smith et al. 2014, Fischer et al. 2014, Atlin et al. 2017, Voss-Fels et al. 2019a). Further, there are concerns that the current rates of genetic improvement for most crops are slower than required to meet global food security targets for this century (Ray et al. 2013, Fischer et al. 2014, Bailey-Serres et al. 2019). Thus, there is always interest in opportunities to accelerate breeding (Atlin et al. 2017). Improved understanding of trait genetic architecture and the physiological understanding of trait contributions to yield potential and adaptation to different environmental conditions are common elements of research efforts that underpin breeding programs (e.g., Jackson et al. 1996, Mace et al. 2013, 2019, Alam et al. 2014a,b, Chenu et al. 2018, Messina et al. 2019, Bailey-Serres et al. 2019, Wu et al. 2019, Wang et al. 2020). Translation of the trait knowledge created from such trait-focused research into accelerated outcomes from breeding for yield potential and yield stability requires integration of the trait G2P models into the operation of the target breeding program (Figure 1; Cooper et al. 2014b, Gaffney et al. 2015, Messina et al. 2018).

Today, with continuing advances in genomic technologies (Morrell et al. 2012, Bevan et al. 2017, Yuan et al. 2017), high throughput trait phenotyping (Araus and Cairns 2014, Araus et al. 2018, van Eeuwijk et al. 2019, Reynolds et al. 2020) and environmental characterisation (Cooper and Hammer 1996, Chapman et al. 2000a,b,c, Chenu et al. 2011, Cooper et al. 2014a,b, Costa-Neto et al. 2020, Resende et al. 2020, de los Campos et al. 2020), there is parallel interest in studying trait genetic variation across scales of biological organization, spanning from genome sequence to the integrated multi-trait phenotypes of crops and their performance in agricultural systems (e.g., Cooper et al. 2002a, 2005, Hammer et al. 2006, 2019, Messina et al. 2011, Marshall-Colon et al. 2017, Chenu et al. 2017, Bailey-Serres et al. 2019, Peng et al. 2020). An important element for the study of trait genetic variation within the RPG of a breeding program (Figure 1) is the relationship between the different alleles of the genes contributing to trait genetic architecture and the effects of the alleles on the trait phenotypes for the individuals possessing the different combinations of the alleles (Figure 2). Throughout, we will refer to this relationship between the combinations of alleles of genes and expression of trait value (phenotype) as the trait G2P link function. Quantitative genetics has developed a diverse range of statistical genetic models to quantify the effects of the alleles of genes on trait phenotypes (Falconer and Mackay 1996, Walsh and Lynch 2018). Recently, there has been interest in exploring novel applications of quantitative genetics for trait-focused plant breeding through utilising and developing the set of functional relationships contained within mechanistic crop models and based on coordinated multi-trait G2P link functions for (Cooper et al. 2002a, Chapman et al. 2003, Cooper et al. 2009, Messina et al. 2011, 2018, Technow et al. 2015, Chenu et al. 2017, 2018, Bustos-Korts et al. 2019a,b). Extensions consider crop improvement through combining breeding and agronomy (Messina et al. 2020, Hammer et al. 2019, Cooper et al. 2020). Throughout, we will refer to these alternative forms of the infinitesimal models that use the hierarchical structure of the crop growth model as CGM-G2P link functions.

**Figure 2.**
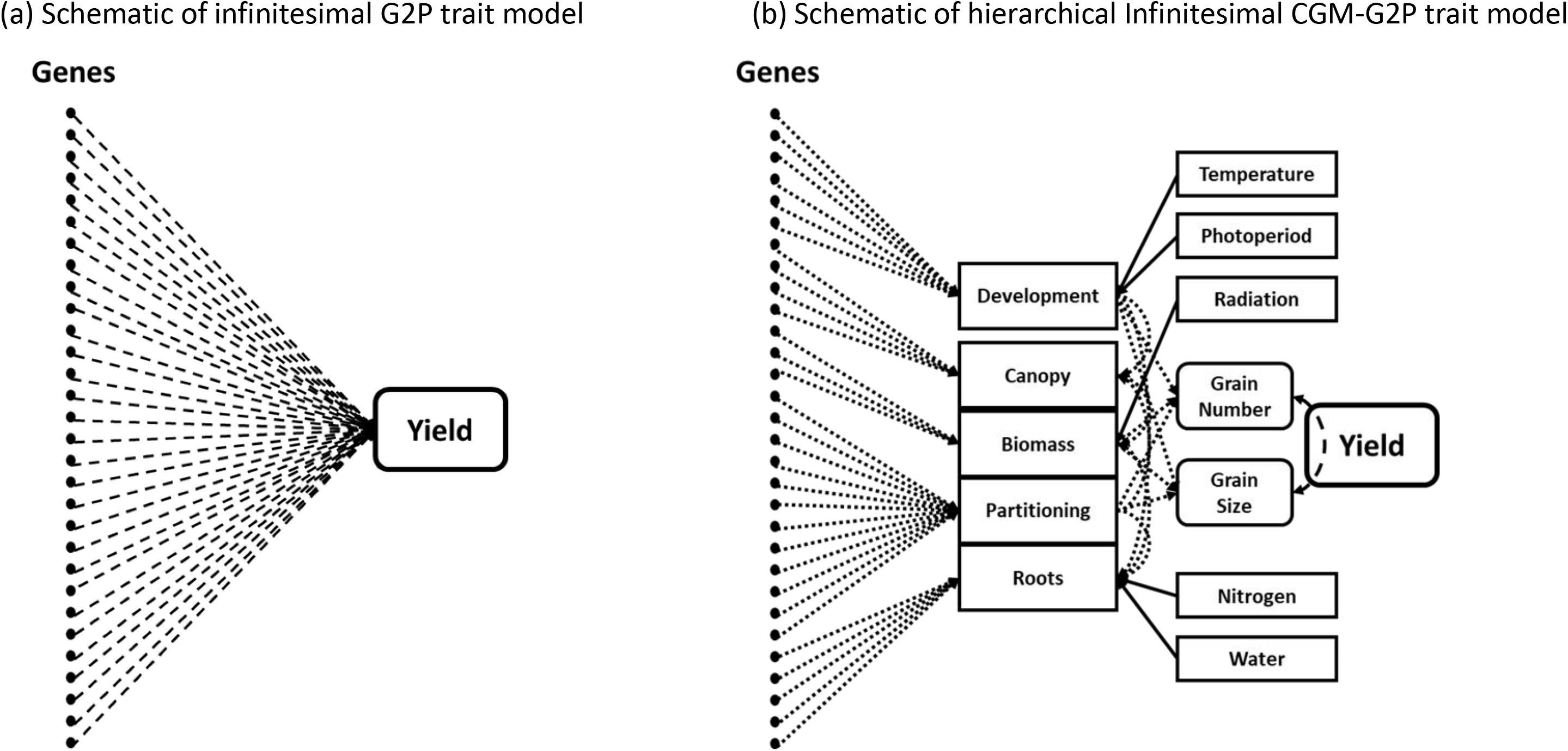
Schematic representation of alternative gene-to-phenotype (G2P) models of trait genetic architecture: (a) Infinitesimal G2P trait model where genes are directly associated with the measured output trait of interest, (b) Hierarchical infinitesimal gene-to-phenotype model (CGM-G2P) using a crop growth model to provide the hierarchical structure where genes are connected to traits at lower levels in the hierarchy to predict traits at higher levels in the CGM hierarchy.

For the purposes of linking trait genetics with crop models we are interested in the connections between our continually evolving mechanistic understanding of plant genomes, trait genetic architecture, the forces that have shaped the standing genetic variation for traits with the RPG of a breeding program and our ambitions to explain trait variation among the cultivars created by plant breeders and tested by agronomists for their performance within a TPE. We discuss some important implications for response to selection of different properties of the trait G2P relationships within crop models, predominantly from the perspective of applied plant breeding programs. We draw from our experience working towards understanding and predicting grain yield variation for cereals (e.g., Bänziger and Cooper 2001, Chapman et al. 2003, Duvick et al. 2004, Messina et al. 2011, 2018, 2019, Cooper et al. 2014a,b). However, the principles are general and can be applied to other traits, crop species and agricultural systems. We aim to draw attention of the crop modelling community to the many opportunities that exist, at the interface between quantitative genetics and mechanistic crop models. Of particular importance is the opportunity to enhance the design of prediction-based methods for crop improvement, and benefit from including contributions from trait genetics into mechanistic crop growth models (Cooper et al. 2005, Voss-Fels et al. 2019b, Messina et al. 2020). While we note there can be a continuum of G2P models, for purposes of illustrating the possible roles of crop models we will contrast two general classes of trait G2P models (Figure 2). The first is representative of the traditional infinitesimal trait G2P models of quantitative genetics, where there is no explicit use of a crop model (Figure 2a). The second, a CGM-G2P, is representative of hierarchical models of trait genetic architecture, where the crop model provides the hierarchical structure of the G2P model and the definition of the traits and their contributions (Figure 2b). It should be understood that there are many ways to construct trait G2P models and to combine these with the hierarchical structure of crop models. The schematics shown in Figure 2 serve to highlight two general approaches, where the key distinction is based on whether the hierarchical structure of a dynamic biological model is not (Figures 2a, 3) or is (Figures 2b, 4) used to connect genome to phenome and quantify gene effects (Cooper et al. 2005, Houle et al. 2010, Barghi et al. 2020).

**Figure 3.**
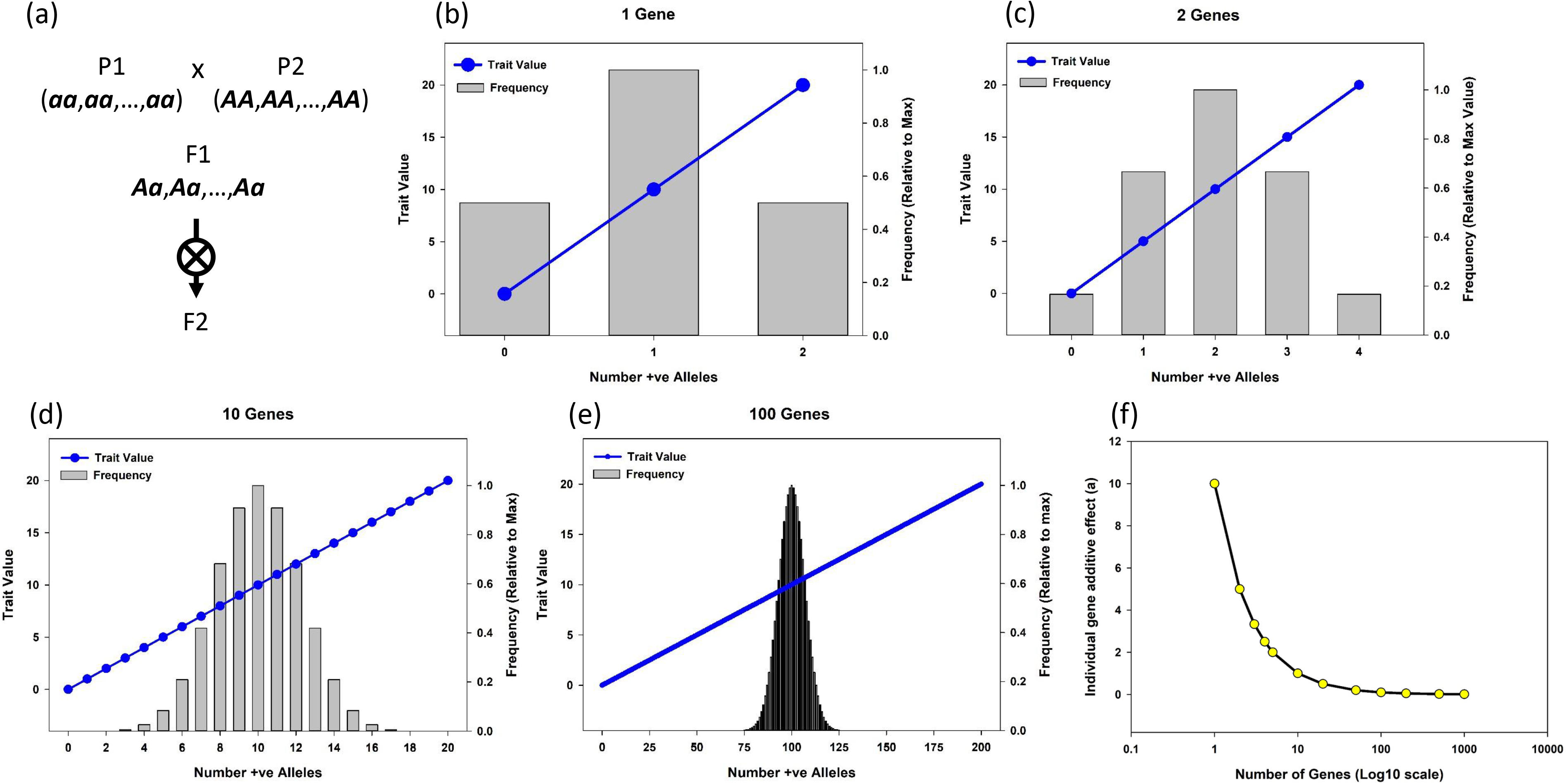
An example of applications of a continuum of additive trait genetic models to an output trait to demonstrate two properties of the infinitesimal model applied within the context of a reference population of genotypes (RPG); (1) the expected frequency distribution of genotypes, where the genotypes are categorised based on the number of alleles that increase trait values summed across all contributing genes, and (2) the trait Gene-to-Phenotype (G2P) link function based on the relationship between the number of alleles that increase trait values, summed across all contributing genes, and the expected trait value of the genotype. (a) The crossing scheme used to create the F2 RPG. (b) Single gene Mendelian model for the total trait genetic variation in the RPG. (c) Two gene model for total trait genetic variation in the RPG. (d) Ten gene model for total trait genetic variation in the RPG. (e) 100 gene model for total trait genetic variation in the RPG. (f) Expected relationship between the number of genes and the size of the effect of each gene for a range of models for total trait genetic variation in the RPG.

**Figure 4.**
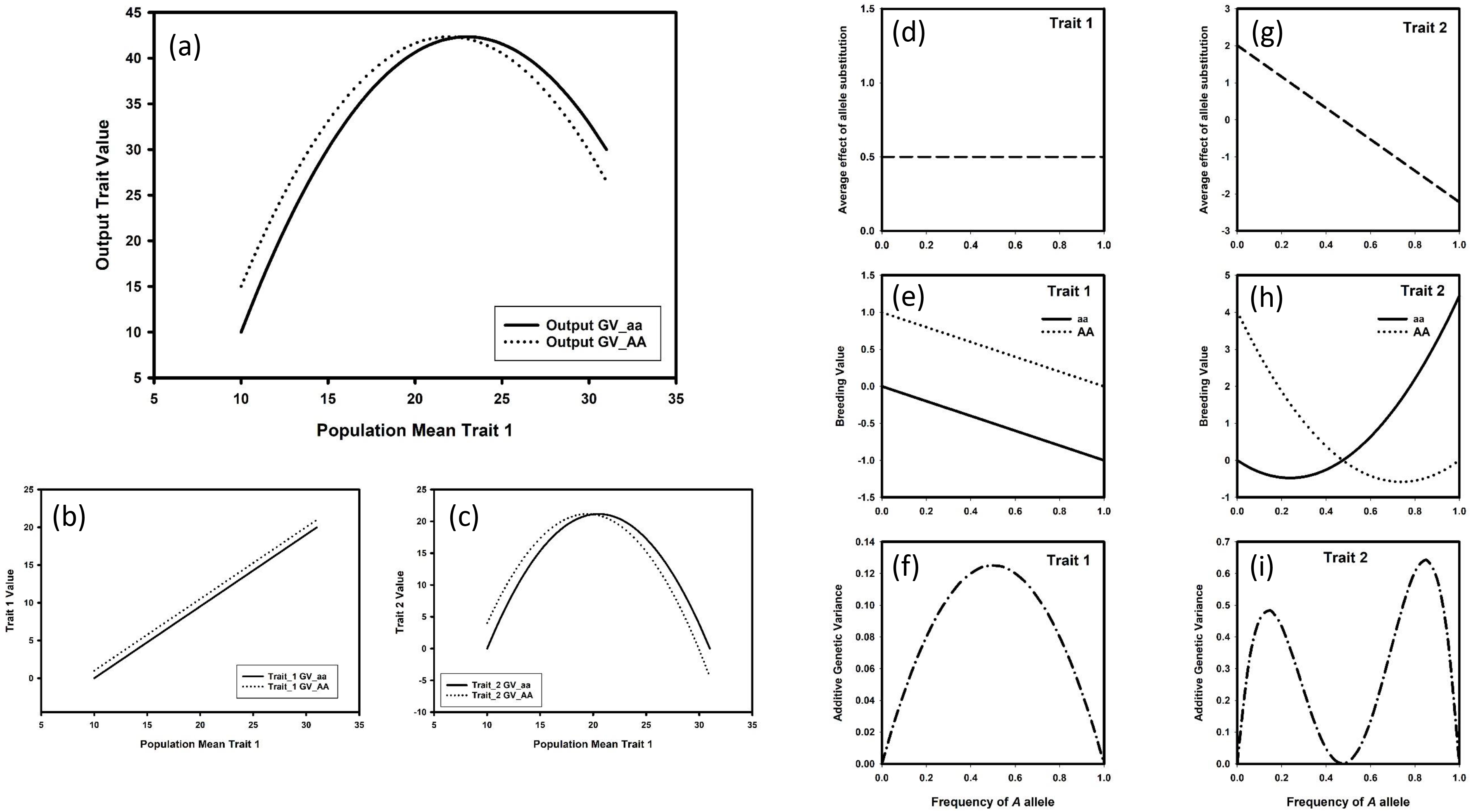
Hypothetical CGM-G2P model to demonstrate a “Gene’s eye view” of the allele effects for a single gene (***Gene_A***) for traits within the context of a hierarchical trait structure. The hypothetical model assumes an output trait (a) at the highest level in the hierarchy is influenced by two traits that both operate at lower levels in the hierarchy of the model; (b) Trait_1 and (c) Trait_2. Sub-figures (a), (b) and (c) focus on the contrast in trait values between genotypes ***AA*** and ***aa*** for ***Gene_A*** as the population mean ranges from a low to high mean value of Trait_1 within the reference population of genotypes (RPG). For the output trait Genotypes ***AA*** and ***aa*** both have a non-linear quadratic relationship with the RPG mean for Trait_1. There is a cross-over of output trait values between genotypes ***AA*** and ***aa***. The responses of genotypes ***AA*** and ***aa*** (a) can be partitioned into a contribution from Trait_1 (b, d, e, f) and Trait_2 (c, g, h, i). For Trait_1 (b) Genotypes ***AA*** and ***aa*** both express a linear increase as the RPG mean value of Trait_1 increases and there is no-crossover between genotypes ***AA*** and ***aa***. Trait_2 (c) is conditioned by Trait_1 and as the RPG mean value for Trait_1 increases Trait_2 has a non-linear quadratic relationship with the RPG mean for Trait_1 and there is a crossover between genotypes ***AA*** and ***aa***. From the G2P relationships depicted for Trait_1 (b) and Trait_2 (c) the gene’s eye view contributions of the alleles of ***Gene_A*** to the components of the breeder’s equation can be determined; Trait_1 (d) average effect of allele substitution, (e) breeding value of genotypes ***AA*** and ***aa***, (f) additive genetic variance; Trait_2 (g) average effect of allele substitution, (h) breeding value of genotypes ***AA*** and ***aa***, (i) additive genetic variance.

It is important to note that within the context of plant breeding, individuals and their genotypes are temporary units within the RPG (Figure 1). Whereas the alleles of the genes represent an enduring basis for studying trait genetics over cycles of breeding and linking genetics to crop growth models. For sexually reproducing crops the alleles of genes, not the genotypes, are transmitted from one generation to the next, from parents to offspring. Therefore, the genes and the alleles persist across generations, not the genotypes. New genotypes are continually constructed from the allele combinations of the genes, following the genetic segregation and recombination that occurs during meiosis, as copies of the genes are transmitted through gametes from parents to offspring. The inclusion of a sexual reproduction step within the cyclical nature of breeding programs (Figure 1b) provides an important motivation for developing a quantitative genetics understanding of traits from the gene and allele level (Falconer and Mackay 1996, Lynch and Walsh 1998, Walsh and Lynch 2018) rather than only at the genotype level. Thus, we need to consider applications of G2P and CGM-G2P models for prediction of the trait phenotype of an individual and the prediction of the individual’s trait breeding value. The former determines the trait performance of the individual and the latter determines the individual’s ability to contribute to the performance new individuals in future cycles of the breeding program. This is an important distinction to draw between gene-to-phenotype and genotype-to-phenotype trait models and their implications as G2P link functions for modelling breeding programs.

For quantitative traits, such as grain yield, the infinitesimal model of quantitative genetics (Falconer and Mackay 1996, Lynch and Walsh 1998, Barton et al. 2017, Walsh and Lynch 2018) underpins many of the genetic principles that are used together with the breeder’s equation to model and predict response to selection by breeding programs. While it is understood that quantitative traits, such as yield, are an outcome of many lower level traits, most breeding programs will focus testing and selection directly on the key endpoint traits. The effects of the alleles of genes on the trait phenotypic values of individuals, based on their genotypes for multiple genes, provides the gene-to-phenotype link function. The framework of the gene-to-phenotype trait model also determines the breeding values of the individuals. Thus, for continuation of the breeding program cycles we are interested in a G2P trait model that enables prediction of the trait values of individuals as potential cultivars as well as the trait breeding values of the individuals.

Physiological investigations provide an important role in developing our understanding of the traits that ultimately determine grain yield of crops and the variation among cultivars within the context of agricultural systems (Evans 1993, Jackson et al. 1996, Cooper and Hammer 1996, Connor et al. 2011, Fischer et al. 2014). However, many investigations to understand the physiological basis of traits and the variation among cultivars for the traits do not take into consideration trait genetic architecture and the implications of the infinitesimal model for genetic improvement over cycles of a breeding program (Figure 1). Physiological investigations to quantify contributions of traits to improved yield performance have focused largely at the level of studying the trait phenotypes for contrasting cultivars, usually with unknown genotypes for the genes that determine the contrasting trait phenotypes of the cultivars. Often the set of cultivars chosen for physiological investigations includes extreme contrasts. Consequently, the genotypic variation for the traits, and their relationships, within the set of cultivars included in experiments focused on understanding trait physiology does not represent the trait variation and relationships within the RPG of the breeding program. Thus, making application within the context of the breeder’s equation difficult. A common outcome from such trait physiological research is the definition of trait ideotypes as proposed breeding targets, without any knowledge of the genetic architecture of the traits or of the genetic changes that are necessary to create the ideotypes within the RPG. Further, the ideotypes rarely are considered within the context of the Target Product Profiles that the breeding programs are designed to create.

There are promising opportunities to improve on this situation by conducting research to link an understanding of trait genetics from such physiological research with mechanistic crop models and to do this within the context of the RPG of the breeding program (e.g. Cooper et al. 2002a, Chapman et al. 2003, Cooper et al. 2014, Chenu et al. 2017, 2018, Messina et al. 2018, Hammer et al. 2019, Bustos-Korts et al. 2019a,b). This would provide a foundation for a wide range of applications of crop models to crop improvement that takes into consideration the processes that are involved in plant breeding to create new genotypes with improved trait performance (Figure 1). Our aim is to highlight potential transdisciplinary pathways for integration of genetic principles within the framework of crop models and to encourage further developments (Hammer et al. 2019).

## G2P Definitions

For the link between quantitative genetics and crop modelling to be a success, terminology needs to be aligned across the contributing scientific domains; genetics, breeding, physiology, agronomy. For discussion of the relevant concepts from quantitative genetics, we follow the terminology of Falconer and Mackay (1996). We then use this terminology in combination with a series of illustrative examples to introduce key genetic principles, beginning with the infinitesimal model of quantitative genetics (Figure 2a, Figure 3). Specialists will seek further elaborations (Falconer and Mackay 1996, Lynch and Walsh 1998, Barton et al. 2017, Walsh and Lynch 2018, Messina et al. 2018).

### Genetics Primer

The genome of an individual is the complete set of chromosomes. Here we restrict our discussion to diploid genomes. Diploid individuals have two copies of the full set of chromosomes. A gene is a segment of DNA with a position on a chromosome, referred to as the gene’s locus. ***Gene_A*** has two alleles ***A*** and ***a***. The alleles differ in some aspect of the base sequence for the DNA segment that defines the gene. Each diploid individual possesses two alleles. The different combinations of the two alleles can produce three different genotypes; the two homozygous genotypes ***aa*** and ***AA*** and the heterozygous genotype ***Aa***. An individual is assumed to have one of the three genotypes for ***Gene_A***. Traits measured on an individual provide a measure of the trait phenotype for the individual. Associations can be established between the trait phenotype and the genotype of an individual if the genotypes of the individuals are known. A Quantitative Trait Locus (QTL) is a region on a chromosome identified through a statistical association between a gene and a trait phenotype. For discussions here, we consider the case where ***QTL_A*** can be associated with ***Gene_A***. The genotype of individuals for ***Gene_A*** can be determined by a range of methods that enable reading of the DNA base sequence for the gene (Rafalski 2002, Palaisa et al. 2004, Edwards and Batley 2010, Morrell et al. 2012, Mace et al. 2013, Shendure et al. 2017, Bevan et al. 2017, Yuan et al. 2017, Bukowski et al. 2018, Wallace et al. 2018, Voss-Fels et al. 2019b, Khan et al. 2020, Jensen et al. 2020). Through appropriate experimental design, a value for a trait can be assigned to each genotype of the gene or QTL, based on the allele combinations, to quantify the average effect of allele substitution (e.g., Falconer and Mackay 1996, Cooper et al. 2005, Buckler et al. 2009, Wang et al. 2020, Boyles et al. 2019). These values can provide a measure of the contribution of a gene or QTL to the total genetic variation and to the trait breeding values of individuals within the RPG.

For demonstration purposes, we consider an F2 generation to introduce quantitative genetic concepts involved in quantifying the effects of alleles of genes contributing to trait variation (Figure 3). The F2 generation is a typical unit of the RPG of a breeding program. Plant breeding programs work with multiple F2 populations simultaneously. Depending on scale, a breeding program may be operated based on tens to thousands of F2 populations to initiate each cycle (Cooper et al. 2014a). The F2 generation is created by first crossing two inbred parent individuals to create an F1 generation. In the current example, the inbred parents were selected to contrast for the phenotypes of the trait of interest and assumed to contrast for the genotypes of all genes controlling the genetic variation for the trait. This is the ideal situation and serves for purposes of discussion. For example, for the case where a single gene controls the trait genetic variation, the genotypes would be Parent 1 ***aa*** and Parent 2 ***AA.*** For the case where two genes control the trait variation, the genotypes would be Parent 1 ***aa***,***aa*** and Parent 2 ***AA***,***AA***; the commas separate the genotypes for the two contributing genes. The model can be extended from single-locus Mendelian genetics to the multi-locus infinitesimal model by considering increasing numbers of genes (Figure 3). The F1 generation, created by crossing the contrasting inbreds, is represented by the heterozygous genotype for the gene that differed between the parents; for one gene the F1 is ***Aa***, for two genes the F1 would be ***Aa***,***Aa***. The F2 generation is then created by self-pollination of the F1 individual; equivalent to crossing the F1 individual with a copy of itself.

## Traits from a Quantitative Genetics Perspective

Quantitative genetics provides a framework for predicting the expected trait values of the genotypes that can be created from the values of the alleles that combine to define the genotypes (Figure 3). The effects of the alleles of genes on the trait genotypic and phenotypic values of individuals provides the gene-to-phenotype link function. We first consider some properties of the foundational finite-locus and infinitesimal G2P models widely used within quantitative genetics (Figure 2a). We then expand on the concept of a CGM-G2P gene-to-phenotype link function to consider crop growth models (Figure 2b).

Falconer and Mackay (1996) provide an introduction to trait quantitative genetics for diploid genomes (Figure 3). Their foundation builds from quantifying the relative effects of two alleles (Nominally defined here as ***A*** and ***a***) for a single locus on trait values for the three possible genotypic combinations of the two alleles (***AA***, ***Aa*** and ***aa***). An ordering of these three genotypes can be defined in terms of the number of one of the alleles that is possessed by each of the three genotypes; e.g., considering the ordering of the genotypes in terms of the number of ***A*** alleles possessed, the genotypes would be ordered as ***aa***, ***Aa*** and ***AA***, with 0, 1 and 2 of the ***A*** alleles, respectively. Using this ordering, a graphical relationship can be constructed between the number of the ***A*** alleles possessed by the genotypes and their trait values. For demonstration purposes, we consider the case where the range of the values for the trait can be partitioned into equally sized components associated with the effects of the alleles for the gene within an F2 RPG (Figure 3). For the example depicted in Figure 3 the trait value range is defined as 20 units of measurement, commencing with a trait value of 0. Thus, for a single gene the three genotypes ***aa***, ***Aa***, ***AA*** have trait values of 0, 10 and 20, respectively, and the additive effect of the ***A*** allele is 10 units (Figure 3b). This graphical view provides a basis for defining the quantitative genetics concept of average effect of allele substitution. For the example depicted in Figure 3b, substituting (replacing) one of the ***a*** alleles for an ***A*** allele in any of the genotypes possessing an ***a*** allele (i.e., genotypes ***aa*** and ***Aa***) results in an increase in trait value of 10 units; e.g. changing ***aa*** to ***Aa*** would increase the trait value by 10 units from 0 to 10, and similarly changing ***Aa*** to ***AA*** would increase the trait value by 10 units from 10 to 20. This framework, applied in combination with the allele frequencies for the RPG, provides the basis for determining the quantitative genetic statistic of average effect of allele substitution. The definition of an “average effect” applies to the RPG. Thus, the “average effect” of all the possible “allele substitutions” within the RPG is dependent on the allele and genotype frequencies within the RPG in combination with the differences in trait values between the genotypes (Figure 3b). The average effect of allele substitution is an important determinant of the additive genetic variation and variation for breeding values within the RPG (Falconer and Mackay 1996). The additive genetic variation is the target of the selection predictions made using different forms of the breeder’s equation, as discussed above (Figure 1).

With a measure of the frequency of the two alleles for the individuals in the RPG for a defined mating structure (Figure 3a), the frequencies of the three genotypes per locus can defined. For the single gene case, the F2 generation represents a RPG comprising of the three genotypes ***aa***, ***Aa***, ***AA***, with expected frequencies of 0.25, 0.5 and 0.25, respectively (Figure 3b). Quantitative genetics provides the framework for generalising these concepts to a wide range of population structures relevant to the RPG of breeding programs.

There are a number of approaches that can be used to extend the views of the link between gene and trait phenotype and frequency distribution of genotypes in the RPG from a single gene to accommodate multiple genes. For demonstration purposes, the approach applied here is to categorise the genotypes over the genes contributing to the trait by summing the number of the ***A*** Alleles that increase the value of the trait over all of the contributing genes. For example, for two genes five genotype categories can be defined based on whether the genotypes have 0, 1, 2, 3 or 4 of the ***A*** alleles over the two QTL. From this categorisation of genotypes in terms of allele numbers, the relationship between the trait value and the number of ***A*** alleles across both genes and the frequency distribution of the five genotype classes within the RPG can be constructed for the five genotypic classes (Figure 3c). Both of these representations of characteristics of the genetic model, the G2P relationship and the frequency of the genotype categories, can be extended for increasing numbers of genes; e.g., 10 genes (Figure 3d), and 100 genes (Figure 3e). With increasing numbers of genes involved in explaining the total genetic variation within the RPG the individual contributions from each gene become progressively smaller, hence the concept of the infinitesimal model (Figure 3f). Thus, we can consider a continuum of quantitative genetic G2P models of trait genetic architecture, ranging from a single Mendelian locus (Figure 3b), with large effect alleles, through a series of intermediate finite locus models that allow for increasing numbers of genes (Figure 3c-e), each with a sequence of decreasing allele effect sizes (Figure 3f), through to the extreme infinitesimal model of quantitative genetics (Walsh and Lynch 2018), with many genes, each making a vanishingly small contribution to the total trait values and the total genetic variance in the RPG.

Importantly, we emphasise that there is a G2P link function that connects the number of alleles possessed by a genotype, across all contributing genes, to the expression of the trait value for the genotypes in the RPG. For the continuum of additive genetic models considered in this example, the G2P link function is linear (Figure 3). Applying the linear G2P link function, within the context of the RPG, the trait values of genotypes can be predicted from the allele configurations of the underlying genes controlling the trait. Thus, if the G2P link function holds for the RPG it is possible to predict the trait values for the genotypes within the context of the RPG, define target genotypes and design breeding strategies to move the RPG towards the target genotypes. The number of cycles of the breeding program required to move the RPG towards the target genotypes (Figure 1) will depend on the genetic architecture of the trait (Figure 3) and the efficiency of the breeding strategy (Cooper et al. 2005). Today, with access to genomic technologies there is considerable interest in using genomic prediction methods to accelerate the rates of trait genetic improvement that can be achieved by breeding programs (Meuwissen et al. 2001, Bernardo and Yu 2007, Cooper et al. 2014a, Voss-Fels et al. 2019b).

From a plant or crop physiology perspective it can be argued that the trait values of the genotypes is of primary interest for prediction of genotype performance for a TPE. Thus, it is possible to connect the trait genetic variation to trait values at the genotype-to-phenotype level, rather than at the gene-to-phenotype level as proposed here. This approach is appropriate where prediction of trait performance for specific genotypes is of direct interest and the numbers of genotypes to be considered is manageable. However, to connect the prediction framework to applications for breeding aimed at creating new genotypes this approach is limited in two respects. Firstly, for cases where the genotypes of interest have yet to be created by the breeding program, direct high-throughput phenotyping is not feasible. Secondly, there is a significant combinatorial challenge when even a modest number of genes control traits. The number of potential genotypes that can be created, and which would need to be phenotyped, rapidly becomes large as the number of genes increases. For a trait where genetic variation within the RPG is controlled by 10 genes, with 2 alleles per gene, the number of potential genotypes is 3^10^ = 5.90×10^4^ (Figure 3d). For 100 genes the number of potential genotypes is 3^100^ = 5.15×10^47^ (Figure 3e).

Clearly, for complex quantitative traits there are much smaller numbers of genes and alleles than the number of potential genotypes. Thus, within the domain of quantitative genetics we seek methods for predicting the trait values of genotypes from measures of the effects of the alleles of the genes.

We can further note that for any sample of individuals taken from the RPG and included in our experiments the target individuals we ultimately seek to create and predict are expected to be outside of the fraction of the total genotype space we can currently observe (e.g., Figure 3e). Therefore, the genetic prediction space where our target genotypes may reside is in all likelihood often outside of the genetic sample space of the data sets that we are currently using to study trait physiology and train the models to be used for genetic prediction. Where the assumption of a linear G2P link function is valid, prediction of the trait values for target genotypes outside of the current genetic sample space is valid (e.g., Figure 3). However, non-linear relationships among traits and between traits and environmental variables is the expectation (Hammer et al. 2006, Messina et al. 2011, 2019, Technow et al. 2015, Wu et al. 2019). Therefore, we require a methodology for dealing with such non-linear relationships. Mechanistic crop models open a range of interesting opportunities to address this shortfall (Figure 2b). Below, we use a non-linear CGM-G2P example to demonstrate applications.

## Applications

### Gene-to-Phenotype (G2P) Modelling for Traits

There is a long history of combining genetic models, using the principles of quantitative genetics, together with computer simulation to investigate dynamic properties of biological systems (Fraser and Burnell 1970, Kempthorne 1988). These early simulation ideas resulted in bespoke software developed to obtain specific answers for specific problems. Some of these studies focused on topics relevant to plant breeding. Success with these early simulation approaches motivated the development of flexible software platforms for modelling a wide range of plant breeding programs (Podlich and Cooper 1998, Cooper et al. 2002b, Li et al. 2012). This has stimulated further interest in developing flexible software tools for a range of applications in plant breeding (e.g., Sun et al. 2011, Faux et al. 2016, Jahufer and Luo 2018, Liu et al. 2019, Gaynor et al. 2020). For most of the software platforms, consideration of the G2P functions for traits has been restricted to the classical quantitative genetic models (Figure 2a; Falconer and Mackay 1996, Walsh and Lynch 1998), with properties similar to those depicted in (Figure 3).

Chapman et al. (2003) introduced an alternative approach for using a mechanistic CGM as a CGM-G2P multi-trait link function, following the schema depicted in Figure 3b. This CGM-G2P framework was enabled by linking the genetic modelling capabilities available within the QU-GENE software (Podlich and Cooper 1998) with the sorghum crop model available in the APSIM platform (McCown et al. 1996, Holzworth et al. 2014). They focused on modelling a sorghum breeding program, following the schema depicted in Figure 1, and demonstrated examples for the objective of improving grain yield for the Australian dryland TPE, which included both drought and favourable (non-drought) environments. The sorghum CGM was used as a multi-trait framework to establish the CGM-G2P link function to quantify QTL effects for grain yield, while no QTL had direct effects on grain yield. The growth and development processes and trait functions included in the CGM determined all QTL effects on grain yield. The QTL were associated with coefficients within the functions for specific traits that were included in the sorghum CGM. The QTL alleles had contrasting effects on the traits and had an effect on grain yield whenever the trait was influential upon grain yield. Technow et al. (2015) used the same hierarchical framework (Figure 2b) to define a CGM-G2P link function for grain yield of maize to enable and demonstrate the advantages of the CGM-WGP genomic prediction methodology for breeding applications. These same approaches have been applied to model maize breeding strategies (Messina et al. 2011, 2018) and to support the development of maize hybrids with improved drought tolerance and yield potential (Cooper et al. 2014b) for the US corn-belt TPE (Gaffney et al. 2015, McFadden et al. 2019).

### The Role of Crop Models in G2P Link Functions

We now turn our attention to considerations for applying the principles of quantitative genetics to the study of trait genetic architecture and trait prediction for breeding (Figure 1) using CGM-G2P hierarchical models (Figure 2b). A primary motivation for considering crop models as G2P link functions for the trait targets of breeding programs is the potential to improve prediction applications for plant breeding and more generally for crop improvement for the complex situations that result in important deviations from the assumptions of linear, additive trait G2P models. Following the framework introduced by Technow et al. (2015), in principle, if we can identify genes controlling traits and through an appropriate CGM-G2P link function, quantify the effects of the alleles of these genes on the expression of trait phenotypic values of plants and their contributions to crop performance in agricultural environments, it is possible to predict the expected trait phenotypic values for the genotypes that can be created by selection within a plant breeding program. To do so we need to establish a common gene-to-phenotype terminology across the disciplines of crop breeding and crop physiology for the development of a suitable crop modelling framework that can incorporate gene-to-phenotype link functions for traits. With such a framework the ability to predict the genotype-to-phenotype relationship for traits at multiple levels in the CGM hierarchy would be an important outcome.

Following the discussion of the breeder’s equation above (Figure 1), to demonstrate some key quantitative genetic principles that will impact application of prediction methods for complex traits we consider the connection of trait genetics to crop models at the gene level. Firstly, we take a “gene’s eye view” of traits through the lens of the CGM. Secondly, we consider some implications of using crop models to enable CGM-G2P link functions for trait prediction applications in plant breeding.

### Gene’s Eye View of trait genetic effects via a Crop Model

Given the combinatorial challenges associated with phenotyping traits for the large number of potential genotypes that can be generated from even a modest number of genes, we have argued that there are important advantages for linking trait genetics to crop growth models through gene-to-phenotype link functions and pursuing further development of such model-based approaches (Figure 2b, Technow et al. 2015). This provides a framework, using the machinery of quantitative genetics, to enable the prediction of genotype trait values from gene and allele effects when a suitable CGM-G2P link function can be defined in terms of the crop model (Messina et al. 2018). Following Wade (2002), we consider a “gene’s eye view” of trait effects using as the CGM-G2P link function the relationships between environmental inputs and traits defined by a crop model.

For the infinitesimal models of trait genetic architecture a distinction is made between the additive and non-additive effects of the alleles of genes and gene combinations (Wade 2002, Cheverud and Routman 1995, Falconer and Mackay 1996, Lynch and Walsh 1998, Walsh and Lynch 2018). A similar distinction can be made for genes influencing traits when using a crop model as the CGM-G2P link function (Figure 2b). Among the genetic phenomena that are commonly identified with non-additivity, two are of direct relevance to linking genetics with crop growth models; genotype-by-environment interactions and epistasis (Hammer et al. 2006, Technow et al. 2015, Messina et al. 2018). Genotype-by-environment interactions refers to changes in the relative trait values of genotypes with changes in environment (Comstock and Moll 1963, Cooper and Hammer 1996). Genotype-by-environment interactions have long been familiar to the developers of crop models and have been targets for investigation using crop models (e.g., Hammer and Vanderlip 1989a,b, Hammer et al. 1989, Chapman et al. 2002). Epistasis refers to gene-by-gene interactions for trait values of genotypes (Falconer and Mackay 1996, Lynch and Walsh 1998, Wolf et al. 2000, Walsh and Lynch 2018). This genetic phenomenon is less familiar to the domains of crop physiology and agronomy. However, some aspects of epistasis have been investigated as trait-by-trait interactions and their implications for selection and prediction investigated using crop models (e.g., Chapman et al. 2003, Messina et al. 2011, 2018, Technow et al. 2015).

While much of the quantitative genetics methodology for studying epistasis and its implications for selection response has developed as a statistical framework, there is recognition of the important distinction between statistical and physiological epistasis (Cheverud and Routman 1995). Statistical epistasis can be considered in terms of the effects of alleles and genes on traits within the context of the RPG of a breeding program while physiological epistasis may be considered in terms of the biological relationships represented by the CGM-G2P link function. The statistical effects of the genes and alleles are dependent on allele and genotype frequencies within the structure of the RPG. Whereas the physiological effects, such as the relationships within the crop model, are not dependent on the structure of the RPG. Therefore, we can develop a crop physiological perspective of both GxE interactions and physiological epistasis within the context of the quantitative relationships defined within a crop model. Further, by linking the effects of genes and alleles to traits through the relationships within the crop model we can quantify the effects of genes through the crop model as a CGM-G2P link function. Therefore, the crop model provides a CGM-G2P framework for tackling components of the important non-linear effects of genes and their interaction with each other and the environment. Thus, the crop model as a CGM-G2P link function provides a mechanism for applying the machinery of quantitative genetics for prediction of expected trait values of new genotypes, that are yet to be created by the breeding program, using the currently observable genotypes within the RPG as the basis for determining the effects of genes and their alleles on the traits. The mathematical relationships between inputs and outputs encoded within the crop model provide the basis for predicting the trait values for the new genotypes for the full range of environments within the TPE. Technow et al. (2015) demonstrated that higher levels of trait predictive accuracy can be achieved using the CGM-G2P link function in a simulated maize example. We propose that the demonstrated CGM-WGP prediction advantages within a breeding program cycle will enhance breeding strategies based on such predictions that aim to move the breeding program over multiple cycles beyond the standing genetic variation that can be sampled to construct the G2P training data sets. Thus, enabling predictions of response to selection that can move germplasm beyond the boundaries of the RPG used to parametrise the CGM-G2P link function.

To demonstrate these connections and prediction opportunities we consider a gene’s eye view (Wade 2002) of trait value through a hypothetical non-linear CGM-G2P link function (Figure 4). For purposes of demonstration, here we consider the simulated output trait values expressed at a higher level in the crop model hierarchy for two genotypes ***AA*** and ***aa*** (Figure 4a) that are determined by the levels of two traits operating at a lower level in the crop model trait hierarchy (Figures 4b,c). This hierarchical trait relationship is motivated by the non-linear relationships between traits reported for a maize crop growth model (Technow et al. 2015, Messina et al. 2018). Following Technow et al. (2015), we consider a hierarchical genetic model, where the genetic control of Trait_1, at the lowest level in the trait hierarchy, is based on a linear additive genetic model (Figure 4b). Trait_2 is in turn conditioned by the level of Trait_1 (Figure 4c). The output trait value at the highest level in the hierarchy (Figure 4a) is obtained as the sum of the values of Trait_1 (Figure 4b) and Trait_2 (Figure 4c), plus a background population mean value. Given these relationships we can follow a “gene’s eye view” of the genetic effects for the alleles of one particular gene, ***Gene_A***, for Trait_1 (Figures 4d,e,f) and Trait_2 (Figures 4g,h,i) as the genetic background of the RPG changes. Here the genetic background change represents a synchronised sweep in the allele frequencies from 0 to 1 for all of the genes other than ***Gene_A*** that are involved in the genetic architecture for Traits 1 and 2. The effect of change in the genetic background is quantified in terms of the population mean of Trait_1, the trait that is positioned lowest in the trait hierarchy of the hypothetical crop model.

For ***Gene_A*** we focus on the two homozygous genotypes; ***AA*** and ***aa*** (Figure 4). For Trait_1 there is a linear relationship between the number of positive alleles for ***Gene_A*** and all of the other background genes and the expression of population mean level of Trait_1 (Figure 4b). Thus, as the frequencies of the alleles contributing a positive effect for Trait_1 increase in the RPG the mean value of Trait_1 in the RPG will increase. For ***Gene_A***, we can estimate the average effect of allele substitution for Trait_1, i.e. effect of replacing an ***a*** allele with an ***A*** allele (Figure 4d), the breeding value of the genotypes ***AA*** and ***aa*** for Trait_1 (Figure 4e), and the contribution of ***Gene_A*** to the total additive genetic variance for Trait_1 (Figure 4f) in the RPG. An important feature of the linear G2P relationship for Trait_1 (Figure 4b) is that the average effect of allele substitution for ***Gene_A*** for Trait_1 is positive and constant (Figure 4d) and the Breeding Value of genotype ***AA*** is always positive until the RPG is fixed for genotype ***AA*** (Figure 4e). Thus, the “genes-eye view” for Trait_1 reveals ***Gene_A*** has genetic effects on Trait_1 values that are consistent with the additive quantitative genetic models for traits depicted in Figure 3.

In contrast with the linear G2P link function for Trait_1 (Figure 4b), for Trait_2, there is a quadratic G2P link function (Figure 4c), with the genotypic value of Trait_2 dependent on the background value of Trait_1. Consequently for Trait_2 a different “gene’s eye view is observed for ***Gene_A***. The G2P link function for Trait_2 is similar to the quadratic relationship between canopy total leaf number and grain yield investigated by Technow et al. (2015) for maize. Many other such non-linear relationships between traits can be highlighted for the relationships defined within crop models (Hammer et al. 2006, 2014, 2020, Messina et al. 2009, 2011, 2019, Wu et al. 2019). For the chosen example, given genotype ***AA*** results in a greater value for Trait_1 than genotype ***aa***, this differential effect on Trait_1 contributes to the values of Trait_2 for the two genotypes (Figure 4c). Consequently, in response to the background value of Trait_1 the quadratic response for Trait_2 for genotype ***AA*** increases to the maximum value for Trait_2 and then decreases ahead of that of genotype ***aa***. Following the same approaches used to examine the genetic effects of ***Gene_A*** for Trait_1, for Trait_2, the genetic effects of ***Gene_A*** can be investigated (Figure 4g,h,i). However, unlike for Trait_1, for Trait_2 there is a change in the sign of the average effect of allele substitution (Figure 4g) and a change in rank of the breeding values of genotypes ***AA*** and ***aa*** (Figure 4h), resulting in a more complex contribution of ***Gene_A*** to the total additive genetic variance for Trait_2 (Figure 4i). Thus, the non-linear shape of the G2P link function for Trait_2 translates into a change in the rank of the value of the alternative alleles of ***Gene_A*** that is conditional on the RPG. Further, this change in the rank of the effects of the alleles for Trait_2 translates into a change in the ranking of the two genotypes ***AA*** and ***aa*** for the output trait at the highest level in the crop model hierarchy (Figure 4a). Thus, the same gene, ***Gene_A***, can have different genetic effects on traits depending on the level of the trait in the crop model hierarchy (e.g., Figure 4; Trait_1, Trait_2 or the output trait). Therefore, correctly linking the gene effects to the traits within the crop model hierarchy allows an appropriate adjustment of the effects of the alleles of genes on trait values of genotypes within the context of the RPG, as required for applications of the breeder’s equation for prediction of trait selection response. Importantly, with knowledge of the non-linear relationship between Traits 1 and 2 and the average effects of allele substitution for the contributing genes (Figures 4d,h), regardless of the number of genes, the trait values for Traits 1 and 2 and the output trait for any of the genotypes that can be created within the RPG can be predicted from the effects of the alleles of the contributing genes. This property of the CGM-G2P link function, which is enabled through connection of trait gene effects to trait relationships within the crop model (Figure 2b), was utilised by Technow et al. (2015) and Messina et al. (2018) to extend the predictive skill of genomic prediction beyond that which was achieved when the infinitesimal model was applied directly to the output trait at the highest level of the crop model hierarchy (Figure 2a). Thus, using the hierarchical structure of the CGM within the G2P link function (Figure 2b) enabled an accounting for the non-additive effects associated with GxE interactions and epistasis of traits for prediction of grain yield of maize genotypes. These important results demonstrate at the genetic level how the CGM-G2P multi-trait link function, based on the hierarchical structure of the CGM (Figure 2b), provides a mechanism for improving predictions for many applications in breeding (Figure 1).

### Implications for Crop Improvement

We have discussed motivations and opportunities to enhance the modelling of plant breeding programs (Figure 1) through incorporation of the hierarchical structure of CGMs within the trait G2P link functions that are used to define trait genetic architecture (Figure 2). The CGM-G2P link function provides new opportunities to improve the predictive accuracy that can be expected from applications of the breeder’s equation for the complex traits that are the targets of breeding programs. Messina et al. (2018) defined and applied a hierarchical Bayesian model to demonstrate advantages of the CGM-G2P framework for CGM-WGP genomic prediction methodology. They demonstrated the advantages using a combination of simulation and empirical maize data sets. The improvements in predictive accuracy were achieved due to the capacity of the relationships within the crop model to provide appropriate adjustments for the average effects of allele substitution that account for GxE interactions and physiological epistasis (e.g., Figure 4). This requires extending the connections of trait genetics to crop growth models to the gene-to-phenotype level. With such a framework, we can highlight three general areas for further consideration:

1. ***Enhanced CGM-G2P models for trait genetics***:

- Extending our understanding and predicting trait values for genotypes where there are significant non-additive genetic effects associated with trait GxE interactions and epistasis.
- Enhanced understanding of the pleiotropic effects of genes on multiple traits. The hierarchical structure of CGMs enables explicit specification of pleiotropic effects of genes within the relationships defined within the CGM.
- Targeting investigations of novel sources of trait genetic variation through mapping to establish associations between polymorphic regions of genomes and traits for prediction within the RPG of breeding programs and through gene engineering and gene editing.
2. ***Improved and novel phenotyping***:

- The incorporation of a CGM within the G2P link function opens up new approaches for precision phenotyping of many traits that are difficult to directly measure on the large numbers of individuals created by breeding programs.
- New phenotyping methods combined with the CGM open up new opportunities for characterisation of the environments of the TPE and targeting breeding methods through design of Multi-Environment Trials to match the key environment-types of the TPE to account for GxE interactions throughout the breeding program cycle.
3. ***Enhanced breeding methodology***:

- Plant breeders are interested in predicting the performance of new genotypes in new environments ahead of the need for extensive and expensive multi-environment testing. The inclusion of a CGM within the G2P trait link function opens up many new opportunities for predicting expected genotype trait values that require predictions beyond the boundaries of the genetic sample space of the training populations that are used for G2P model building and where the additive assumptions of the infinitesimal model do not hold. Important applications include predicting strategies for crop genetic improvement for future environments impacted by the anthropogenic effects of climate change.
- Optimal breeding strategies require the application of selection to multiple traits to make progress towards the target Trait Product Profiles. The design of optimal multi-trait selection indices is difficult. CGMs open opportunities for new ways to manage multi-trait selection and design effective multi-trait selection indices for many breeding applications.

## Discussion

The introduction to the key elements involved in modelling a breeding program (Figure 1) provides a focus for linking trait genetics with crop models (Figure 2). The successful link between trait genetics and crop models opens a pathway for developing prediction methods that can increase the scale of crop improvement programs by orders of magnitude (Cooper et al. 2014). We argue that prediction of the expected trait values of genotypes based on the effects of the alleles of the genes through an appropriate CGM-G2P gene-to-phenotype link function, rather than focusing on a genotype-to-phenotype link function, has advantages for breeding applications. Applying the CGM-G2P framework, Chapman et al. (2003) demonstrated the modelling of a sorghum breeding program using the hierarchical structure of a sorghum crop model to account for non-linear relationships among traits and the differential contributions of traits to grain yield for contrasting environments. Applying the same approach, Technow et al. (2015) demonstrated how genomic prediction for grain yield was improved by incorporating a crop growth model into the prediction framework. Combining these approaches provides the foundation for new approaches to model plant breeding programs that enhance our capability to take into consideration the non-stationary effects that genes can have on trait breeding values and the emergent trait phenotypes of individuals within and across cycles of the breeding program.

For the purposes of this review, we have focused on plant breeding that involves a sexual reproduction step between generations of the RPG and therefore, between cycles of a breeding program (Figure 1). For such sexually reproducing plant species, the alleles of genes are passed from one generation to the next with genotypes created anew each generation. Therefore, to predict genetic improvement by plant breeding, when sexual reproduction is involved, it is necessary to connect the genetics of traits to the phenotype at the level of the allele effects rather than only at the genotype level. Thus, to study the outcomes of selection for traits in plant breeding the G2P link function needs to be defined for the gene-to-phenotype level rather than only at the genotype-to-phenotype level. Applying the methodology of quantitative genetics to the hierarchical structure of an appropriate CGM it is, therefore, possible to predict the expected trait values for the genotypes of individuals from CGM-G2P link functions defined at the gene-to-phenotype level (Technow et al. 2015, Messina et al. 2018). We have demonstrated that including a suitable CGM within the G2P link function defined at the gene-to-phenotype levels (Figure 2b) can improve the range of applications for translation of our understanding of trait genetic architecture into the predicted performance profile of genotypes for the range of environments in a TPE.

Successful crop performance in agricultural environments is a consequence of the combined contributions from multiple traits. However, much of our trait research focus has been on one or a few traits. We have struggled to integrate the effects of individual trait studies to successfully predict strategies to improve crop performance for the many diverse environmental conditions encountered within a TPE. In the last three decades, quantitative genetics methodology has been widely applied to map the genetic architecture of plant traits. We have learnt a lot about the genetic architecture of traits and the standing genetic variation for traits within the RPG of breeding programs. However, in most cases we do not know, and are unlikely ever to know, the complete set of genes controlling the standing genetic variation for traits in any RPG. Despite this, with access to a large number of molecular markers distributed across the genomes of crops and ability to measure trait phenotypes of interest, we can map the regions of the genomes where the genes and associated influential sequence polymorphisms are located. For most crops, we now have a toolkit to map and study the effects of genes and QTL for any trait that can be phenotyped in a suitable RPG (e.g., Boer et al. 2007, Buckler et al. 2009, van Eeuwijk et al. 2019, Wang et al. 2020). This mapping foundation has resulted several proposals for prediction-based approaches for breeding based on the traits studied by plant and crop physiologists. However, to date these prediction-based approaches have predominantly remained as research undertakings, and their wide adoption in applied breeding has remained elusive. The current proposals for tackling molecular-assisted breeding by targeting one or a few traits may be limiting the opportunities to apply our growing understanding of trait genetic architecture (Bailey-Serres et al. 2019). Crop growth models provide a quantitative framework, based on well-researched principles of plant and crop physiology, for investigating the contributions of multiple traits to crop performance across a range of agricultural environments (Hammer et al. 2019). Therefore, it now seems appropriate and timely to more fully explore the potential roles of CGMs as suitable gene-to-phenotype (CGM-G2P) link functions for many traits of interest to extend the current investigations of trait genetics. The greater exploration of CGM-G2P link functions would enable extensions of quantitative genetics methodology to study the genetic architecture of crop adaptation strategies that are based on the integrated contributions of multiple traits. The framework presented here applied to CGM-G2P multi-trait link functions will enable molecular-assisted breeding for complex breeding objectives and is therefore relevant across the crop modelling community.

## Acknowledgements

Contribution supported by the Australian Research Council Centre of Excellence for Plant Success in Nature and Agriculture (CE200100015) and the Australian Grains Research and Development Corporation project UOQ1903-008RTX.

